# Fear memory in humans is consolidated over time independent of sleep

**DOI:** 10.1101/2022.03.30.486375

**Authors:** Yuri G. Pavlov, Nadezhda V. Pavlova, Susanne Diekelmann, Boris Kotchoubey

## Abstract

Fear memories can be altered after acquisition by processes such as fear memory consolidation or fear extinction, even without further exposure to the fear-eliciting stimuli, but factors contributing to these processes are not well understood. Sleep is known to consolidate, strengthen and change newly acquired declarative and procedural memories. However, evidence on the role of time and sleep in the consolidation of fear memories is inconclusive. Here, we used highly sensitive electrophysiological measures to examine the development of fear-conditioned responses over time and sleep in humans. We assessed event-related brain potentials (ERP) in 18 healthy young individuals during fear conditioning before and after a 2-hour afternoon nap or a corresponding wake interval in a counterbalanced within-subject design. The procedure involved pairing a neutral tone (CS+) with a highly unpleasant sound (US). As a control, another neutral tone (CS-) was paired with a neutral sound. Fear responses were examined before the interval during a habituation phase and an acquisition phase as well as after the interval during an extinction phase and a re-acquisition phase. Differential fear conditioning during acquisition was evidenced by a more negative slow ERP component (stimulus-preceding negativity) developing before the unconditioned stimulus (loud noise). This differential fear response was even stronger after the interval during re-acquisition compared to initial acquisition, but this effect was similarly pronounced after sleep and wakefulness. These findings suggest that fear memories are consolidated over time, with this effect being independent of intervening sleep.

## Introduction

Fear-related disorders, such as panic disorder, specific phobias, and posttraumatic stress disorder (PTSD) are characterized by pathological fear responses as a result of exposure to strong fear-evoking events. The mechanism underlying the formation of many fear memories is a process of fear conditioning. Fear conditioning is a type of associative learning where an initially neutral stimulus (e.g., sound) – conditioned stimulus (CS) – is repeatedly paired with an aversive stimulus (e.g., electric shock or noise burst) – unconditioned stimulus (US). After multiple pairings, the CS starts eliciting the conditioned response, a physiological and/or behavioral reaction similar to the one observed previously to the US. In differential fear conditioning paradigms, one CS (usually designated as CS-) typically represents a safety signal that is never paired with the aversive US, while another CS (CS+) is predictive of an aversive event. Extinction learning is achieved by repeated presentation of the CS+ without the US, leading to attenuation of the conditioned response, possibly due to a new association, competing with the original fear memory (Milad & Quirk, 2012). Extinction learning is considered a central mechanism for changing maladaptive fear memories in the therapeutic process, e.g., as a part of exposure therapy. Understanding how different factors affect fear memory formation, consolidation and extinction is essential for the advancement of targeted therapeutic interventions for the treatment of anxiety disorders.

Sleep plays an essential role in memory formation, particularly in the consolidation of new memories, i.e., strengthening and stabilization of memory traces after initial acquisition. Sleep facilitates the consolidation of memories in the domains of declarative memory (the memory for facts and personal experiences), and procedural memory (memory for skills such as riding a bicycle) (Diekelmann & Born, 2010; Schimke et al., 2021; Schmid et al., 2020). For emotionally charged memories, the role of sleep in memory consolidation is less clear. Two recent meta-analyses showed no overall effect for preferential sleep-dependent consolidation of emotional over neutral material (Lipinska et al., 2019; Schäfer et al., 2020), and studies on sleep-related consolidation of conditioned fear memories in humans have yielded conflicting results. Some studies found stronger conditioned fear responses after sleep (Menz et al., 2013), whereas others observed no difference between sleep and a period of wakefulness (Davidson et al., 2016; Menz et al., 2016; Zenses et al., 2020) or even stronger differential fear responses after wakefulness (Davidson et al., 2018; Lerner et al., 2021). Thus, the question of whether and how sleep affects the consolidation of fear memories remains largely open.

The research on fear memories has employed a number of different measures to characterize fear. Most studies assessed behavioral and/or peripheral physiology measures of freezing, avoidance, and hyperarousal, which, however, do not necessarily correlate with the mental state of fear (LeDoux & Hofmann, 2018; Mobbs et al., 2019). For example, skin conductance response (SCR), which is the most commonly used peripheral measure of fear conditioning, can be interpreted to manifest expectancy (Gazendam & Kindt, 2012; Luck & Lipp, 2016; Soeter & Kindt, 2010), emotional response (Chen et al., 2021; Wood et al., 2014), elevated arousal (Beckers et al., 2013; Williams et al., 2001), or a mixture of these. On the other hand, the pure brain mechanisms directly related to mental states are less frequently investigated. In humans, neural indices of fear learning have been studied by means of fMRI (Fullana et al., 2016) and EEG/MEG (Miskovic & Keil, 2012). As an advantage, EEG can track fast-changing cortical responses and the event-related potential (ERP) components can help to differentiate different stages of fear processing. The earlier ERP components, such as P1, N1, and P2, can indicate changes in sensory processing of conditioned stimuli with larger amplitudes possibly indicating elevated saliency of threatful CS+ (Miskovic & Keil, 2012), while P3 may represent involuntary attention capture by the CS+ (Pavlov & Kotchoubey, 2019). Late positive potential (LPP), i.e., a slow ERP component that typically starts about 400 ms after CS presentation with a maximum at the posterior electrodes, may be related to attention allocation to emotionally salient information (Schupp et al., 2006), while the stimulus preceding negativity (SPN), i.e., a slow ERP component that typically occurs between 200 and 500 ms before expected relevant stimuli, may indicate affective anticipation of the US, with a more negative amplitude possibly reflecting elevated outcome anticipation (Baas et al., 2002; Böcker et al., 2001; van Boxtel & Böcker, 2004).

Previous fear conditioning research employing EEG identified several ERP components that may serve as indices of conditioned responses. ERP components such as P1 (Pizzagalli et al., 2003), N1 or P2 (Kluge et al., 2011), and P3 (Kotchoubey & Pavlov, 2017; Pavlov & Kotchoubey, 2019; Rothemund et al., 2012), the LPP (Bacigalupo & Luck, 2018; Panitz et al., 2015; Pavlov & Kotchoubey, 2019; Sperl et al., 2021), and the SPN (Baas et al., 2002; Böcker et al., 2001; Ferreira de Sá et al., 2019; Hellwig et al., 2008; Regan & Howard, 1995), have been reported to produce a differential response in the contrast between conditioned stimuli (CS+) and safety signals (CS-). The processes manifested in these different ERP components can help to unveil the mechanisms underlying the consolidation of fear memories.

In the current study, we examined the effects of a post-learning retention interval of sleep or wakefulness on the consolidation of fear memories, assessed with behavioral measures (subjective ratings) and neural correlates (ERP) of fear-conditioned responses. In a within-subject design, subjects acquired auditory-conditioned fear memory and were tested on extinction and re-acquisition following a 2-hour afternoon nap or a respective wake period. We hypothesized that (1) a reliable correlate of classical fear conditioning is observed in ERP components, (2) the retention interval results in fear memory consolidation that manifests itself in stronger conditioned responses (expressed in ERP and, possibly, in subjective ratings) after compared to before the retention interval (i.e., time effect), and (3) fear memory consolidation is further enhanced by sleep, which is evidenced by larger ERP responses (and potentially higher subjective ratings) to fear conditioned sounds after sleep compared to wakefulness during extinction and reacquisition (i.e., sleep effect).

## Methods and Materials

### Participants

The study included 18 participants (8 females) with an average age of 24.7 ± 3.18 (mean±SD) years. All but two participants were students at the University of Tübingen. The inclusion criteria were age between 18 and 40 and being a native German speaker. Exclusion criteria were any neurological or psychiatric disease in the past, taking any medication during the investigation, and smoking. All participants but one were right-handed. Participants gave their informed written consent and were paid for their participation in the study. The study was approved by the local Ethics Committee of the Medical Faculty of the University of Tübingen.

### Procedure and task

The general design of the experiment is depicted in Figure 1a. Each participant visited the laboratory for a total of three afternoons. During the first “adaptation” day, participants had a two-hour nap to adapt to sleeping in the lab environment. Separated by at least 13 days (mean±SD, 27.9±16.8), the second and third days then took place, which consisted of a “sleep” and a “wake” day, respectively. The order of these two days was counterbalanced between participants (no order effects on the EEG and subjective ratings were found). The tests performed on these days were identical; the only difference was the intervention: on sleep days, participants went to bed and had a 2-hour sleep opportunity between the conditioning sessions. On wake days, participants stayed awake for the same time sitting in a comfortable chair whilst changing activities every 30 min between watching a silent nature film and playing a computer game (“bubble shooter”). After both interventions, to give them time to fully awake after the nap, the participants spent additional 30 minutes filling out questionnaires unrelated to the purpose of the experiment. On both days, participants arrived at 1:00 pm at the lab. Before and after the sleep/wake period the participants underwent a fear conditioning paradigm. For the duration of this paradigm, they were asked to stay awake, sit still, and attentively listen to the stimuli with closed eyes; they were not required to perform any actions (i.e., a passive auditory oddball paradigm). To facilitate the probability of falling asleep on the sleep day, before each of the three days, participants were asked to wake up 1 hour earlier than usual.

**Figure 1.**
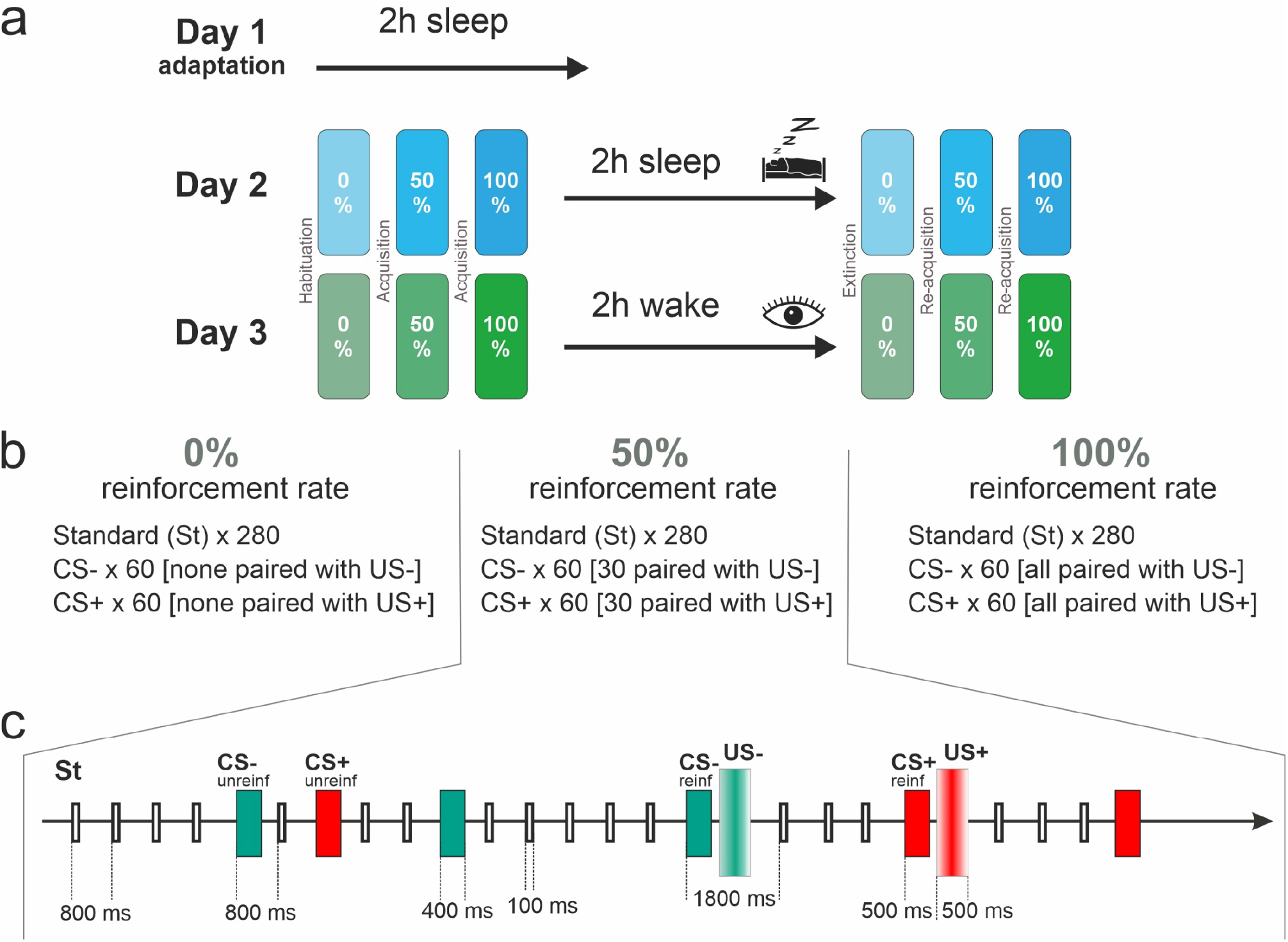
(a) General design. The participants came to the lab three times on three separate days. On the first (adaptation) day the participants went to bed and had a 2-hours sleep opportunity. On the second and third days, the participants either went to bed for 2 hours or remained awake. Before and after the period of sleep or wakefulness the participants underwent a fear conditioning experiment with three blocks of trials. (b) Content of the fear conditioning experiment. 0, 50, and 100% reinforcement rate blocks contained 0, 30, and 60 reinforced conditioned stimuli, respectively, and 280 Standard stimuli. (c) An exemplary sequence of several trials corresponding to a 50% reinforcement rate block with all stimulus types shown. St – standard stimulus, CS+ and CS-– conditioned stimuli, US+ and US-– unconditioned stimuli (US+ – loud noise burst, US- – mildly pleasant sound), reinf – reinforced, unreinf – unreinforced.

All the stimuli were presented auditorily via pneumatic earphones (3M E-A-RTONE). One of the stimuli (Standard) consisted of five frequencies with the fundamental frequency (f1) randomly chosen between 320 and 500 Hz, and f2 = f1*2, f3 = f2*2, f4 = f3*2, and f5 = f4*2. The other two tones were referred to as CS+ and CS-. One of those tones had the main frequency 25% higher, and the other one 25% lower than the fundamental frequency of the standard tone. The frequency assignment was random. The timbre of the CS tones also varied between experimental days including either three or five harmonics per tone. One of the CS tones, randomly selected, was presented to the left ear and the other one was presented to the right ear while the standards were presented to both ears. The duration of the standard tone and the CS tones was 100 ms and 400 ms respectively, with 5 ms fade-in and fade-out. The loudness of the CS tones was 70 dB sound pressure level (SPL), and for the standards it was 65 dB SPL. On each of the two experimental days, the stimuli were generated anew assuring no repetition of the previous day’s stimuli but remained the same within the day.

The CS tones were paired with either an aversive unconditioned stimulus (US+) or a neutral sound (US-). The stimulus-onset-asynchrony (SOA) between CS and US was fixed to 500 ms. SOA for unpaired CS was 800±50 ms (i.e., 750-850 range) and SOA for US was 1300±50 ms (see Figure 1c). The aversive stimulus (US+) was a 1000 Hz sine wave embedded into a burst of white noise (92 dB, 500 ms). The US- was a mildly pleasant sound that consisted of mixed frequencies (70 dB, 500 ms) (the stimuli are available on OSF: https://osf.io/xph69/). Subjective ratings of valence and arousal (see below for description) confirmed that the US+ was indeed perceived as more aversive than the US-, i.e., more arousing (US+_arousal_: 6.31±1.55, US-_arousal_: 3.54±1.89; mean±SD) and more negatively valenced (US+_valence_: 2.40±1.10, US-_valence_: 7.19±1.41; mean±SD). US-served as a neutral control stimulus that allowed us to track sensory responses to the stimulus in a time-locked fashion. For comparison, similarly to ratings for CS+ and CS-after the habituation block (see Results), valence ratings for the standard stimulus were slightly positive (compared to the absolute neutral rating of 5) on average (Standard_valence_: 6.01±1.12, Standard_arousal_: 2.42±1.25; mean±SD). Although mildly pleasant, the US-cannot be interpreted as an appetitive stimulus because alone it did not elicit an unconditioned appetitive response, and no successful conditioning of CS- (i.e., no shift of valence towards more positive values after acquisition) was observed.

In each fear conditioning session (before and after the sleep/wake interval), participants were presented with three blocks of 400 stimuli each, where the rare CSs were interspersed with frequent standard stimuli (Figure 1c). In each block, 280 Standard, 60 CS+ and 60 CS-stimuli were presented with different reinforcement rates: 0%, 50% and 100% (see Figure 1b). In the 0% reinforcement rate block, no US was delivered. In the 50% reinforcement rate block, half of the CS+ and CS-were paired with the respective US. In the 100% reinforcement rate block, all CS+ and CS-were paired with the respective US. All participants had a short self-paced break after each block. The same three blocks with the same stimuli were presented before and after the sleep/wake interval.

The six blocks were named according to the most elaborated and general notation of fear conditioning studies, developed by Lonsdorf et al. (2017). According to this scheme, the 1st block before the sleep/wake interval was referred to as *habituation*, the 2^nd^ and 3rd blocks before the sleep/wake interval, as *acquisition*, (in which the 2^nd^ block employed partial reinforcement), the first block after the sleep/wake interval was regarded as *delayed extinction* (for brevity, just *extinction*, because there was no non-delayed extinction in our experiment), and the last two blocks after the sleep/wake interval, as *re-acquisition*.

### Subjective arousal and valence ratings

Subjective arousal and valence ratings of the stimuli were assessed with 9-point self-assessment manikins to evaluate emotional responses towards the stimuli during the conditioning procedures (Bradley & Lang, 1994). Before the conditioning, for training, the US+ (loud noise) was presented three times, and participants were asked to rate the sound on arousal and valence scales. Immediately after each block of habituation, acquisition, extinction, and re-acquisition each of the stimuli presented in that block was played individually once again (e.g., after the first block only Standard, CS+, and CS-were presented, but after the second and third blocks US+, US-were presented as well), and participants were asked to rate them for arousal and valence.

### Vigilance

The Stanford sleepiness scale was used to assess the participants’ subjective sleepiness (Hoddes et al., 1972). The participants filled out the questionnaire 4 times during each experimental day (before and after each conditioning session of 3 blocks).

For an objective assessment of vigilance, participants were asked to perform a psychomotor vigilance task (PVT) 2 times during each experimental day (before the first conditioning session and before the second conditioning session, i.e., 30 minutes after the end of the sleep/wake interval). In this vigilance test, the participants were asked to place left and right index fingers on two selected buttons of a computer keyboard. A red circle repetitively appeared on either the left or on the right side of a dark screen every 2, 4, 6, 8, or 10 seconds. Participants were asked to press the corresponding left or right button as fast as possible. The task lasted 5 minutes. Median reaction times entered the analysis.

### Sleep duration and quality before the experiment

As a part of the entry questionnaire, participants were asked to estimate their regular sleep duration. The SF-A/R (Schlaffragebogen, "sleep questionnaire”-A, revised) (Görtelmeyer, 2011) was used to evaluate the participants’ sleep quality the night before each experimental day and also following the 2-hour nap on the experimental sleep day. Participants completed the SF-A/R regarding the previous night before the conditioning session.

### Electroencephalography and event-related potential analysis

A 32-channel EEG system with active electrodes (ActiCHamp amplifier and actiCAP slim electrodes, Brain Products) was used for the recording. The electrodes were placed according to the 10-20 system with Cz channel as the online reference and Fpz as the ground electrode. The level of impedance was maintained below 25 kOm. The sampling rate was 1000 Hz.

EEGLAB (Delorme & Makeig, 2004) was used for data preprocessing. Each recording was filtered by applying 0.1 Hz high-pass and 45 Hz low-pass filters (pop_eegfiltnew function in EEGLAB). Then, an Independent Component Analysis (ICA) was performed using the AMICA algorithm (Palmer et al., 2012). As a preliminary step for improving ICA decomposition, a high-pass 1 Hz filter was applied. Application of the low-pass filter could not negatively affect the result of ICA because (1) all ERP components that were of interest in the present study are of much lower frequency than the pass band of the amplifier (specifically, all of them lie well below 20 Hz), and, (2) due to the comfortable position with eyes closed, there was little muscle activity in the signal. It has been shown that removing irrelevant activity from the EEG before ICA can help to improve signal decomposition (e.g., Castellanos & Makarov, 2006; Mannan et al., 2016; Zakeri et al., 2014).

The results of the ICA on the 1Hz filtered data were then imported to the data filtered with 0.1 high-pass filter. Components clearly related to eye movements and high amplitude muscle activity were removed. Additionally, components that were mapped onto one electrode and could be clearly distinguished from EEG signals were subtracted from the data. After this, data were re-referenced to average reference and epoched into [-200 800 ms] intervals with 0 representing CS and US onset, where the [-200 0] interval was used for baseline correction. Epochs still containing artifacts were visually identified and discarded. Finally, before entering the statistical analysis data were re-referenced to averaged mastoids.

We first aimed to test the classic auditory ERPs normally appearing in the auditory oddball paradigm (Barry et al., 2007; Justen & Herbert, 2018; Kotchoubey & Pavlov, 2019). For this analysis, mean amplitudes of the N1, P2, P3a, P3b, and N3 were computed in time windows of 70-125, 140-180, 180-290, 290-360, and 360-500 ms post-stimulus, respectively. We did not directly adopt time windows from the literature because the peak of the components could be affected by conditioning, intensity of the stimuli, and minor changes in the recording environment. Therefore, the time windows were chosen based on group averages collapsed across all experimental conditions and relevant channels (i.e., most commonly used in ERP literature channels reliably reflecting the topographical distribution of the components while being minimally affected by artifacts: Fz, Cz, Pz^1^) – the flatten average approach (Bowman et al., 2020). Specifically, before conducting any statistical analyses, we identified peaks on the flatten average and selected non-overlapping time windows (see Figure S1 for the flatten average). The time windows selection was guided by the number of components typically observed and analyzed in the paradigm (i.e., five: N1, P2, P3a, P3b, and N3, e.g., see Pavlov & Kotchoubey, 2019), peak location in time (on the flatten average), sharpness of the peak (e.g., because the P2 is the fastest and the N3 is the slowest component, the duration of the time windows was shorter and longer, respectively) and the duration of the trial (i.e., 500 ms – the last time window end was fixed at 500 ms).

For the main analyses of the effect of conditioning, time, and type of the intervention, we used SPN time window (180-500 ms) averaged over C3, C4, and Cz channels (for more details, see Results, subsection “Differential conditioned responses in ERPs”).

For estimation of the unconditioned response, we used the N1-P2 amplitude difference as a measure of primary sensory cortical response with an N1 time window of 590-650 ms and a P2 time window of 710-770 ms after CS onset (defined as 60 ms time window around the peaks in the flatten average), averaged over C3, C4, Cz channels. To compensate for a possible drift in the baseline due to the SPN, we used the difference score instead of the ERP amplitude in N1 and P2 separately.

### Polysomnography

To assure that participants were asleep on the sleep day and awake on the wake day in the respective interval, in addition to EEG, chin electromyography (EMG) and electrooculography (EOG, positioned 1 cm lateral and below to the outer canthi of the left eye, and 1 cm lateral and above the outer canthi of the right eye) were assessed.

Before sleep scoring, data were referenced to average mastoids. Sleep scoring was performed visually on 30-s epochs according to standard criteria of the AASM (Iber et al., 2007). The channels F3, C3, and O1 were used for the analysis unless additional information was required to classify an epoch in which case other channels were used in addition.

### Statistics

Most repeated-measures ANOVAs, unless specified otherwise, involved a factor Stimulus (2 levels: CS+ and CS-), Block (3 levels: 0%, 50%, 100% reinforcement rate), Intervention (2 levels: sleep and wake), and PrePost (2 levels: before and after the intervention). The Greenhouse-Geisser correction was applied in the cases of violation of sphericity. Where appropriate, the significant effects and interactions were followed by post-hoc t-tests with Holm’s correction for multiple comparisons. Cohen’s d for paired samples t-tests (dz) is reported as a measure of effect size.

The main results of ERP analyses were additionally confirmed via application of non-parametric cluster-based permutation tests as implemented in Fieldtrip toolbox (Oostenveld et al., 2011). We applied the tests throughout the epoch (0-800 ms) with at least 2 channels threshold for forming a cluster.

Because in several similar fear conditioning studies direct CS+/CS-comparisons were performed instead of a general ANOVA, we also carried out such tests even when they were not indicated by significant ANOVA interactions. As the results were always the same as in our main analysis, these additional analyses will not be reported.

We did not conduct an a priori power analysis and the study was not preregistered. A sensitivity analysis conducted in G*Power (Faul et al., 2007) for the paired t-test (main effect of intervention), the study with N = 18 could detect large effects of dz = 0.71 with 80% power and alpha level of 0.05. A similar analysis for two independent groups (i.e., intervention as a between-subject factor) and the same effect size would require N = 66 (i.e., 33 per group).

The complete statistical results of the main analyses and additional analyses are reported in Supplementary materials.

All statistical analyses were performed with R version 4.1.0 (R Core Team, 2013).

## Results

### Sleep and vigilance

Participants’ average reported sleep duration was 8±0.67 (mean±SD) hours. Sleep duration during the nights preceding the experiment was on average 7.5±0.75 (mean±SD) hours. Sleep quality (scale range 1–5) on the nights preceding the experiment was within the normal range with 4.13±0.28 (mean±SD) on average. All participants successfully fell asleep during the sleep intervention while none of the participants showed any signs of sleep during the period of wakefulness (Table 1). The correlations between sleep duration and differential conditioned responses are reported in Table S1.

**Table 1.** Results of polysomnography on the sleep day.

| Variable | Mean (SD) | minimum | maximum |
| --- | --- | --- | --- |
| N1, min | 18.6 (13.9) | 3.5 | 65.5 |
| N2, min | 43 (17.4) | 6.5 | 65.5 |
| N3, min | 19.9 (16) | 0 | 49 |
| REM, min | 9.9 (11.5) | 0 | 33 |
| N1, % | 22.7 (19.9) | 3.3 | 91 |
| N2, % | 46.6 (15.1) | 9 | 66.7 |
| N3, % | 21.1 (18) | 0 | 62.9 |
| REM, % | 9.4 (10.6) | 0 | 29.3 |
| Sleep latency, min | 9.7 (7.2) | 3 | 35 |
| Total sleep time, min | 91.6 (21.6) | 25.5 | 112.5 |

N1 – stage 1 sleep, N2 – stage 2 sleep, N3 – slow wave sleep, REM – rapid eye movement sleep

Subjective sleepiness measured by the Stanford Sleepiness Scale decreased after the 2-hour nap (mean±SD, before: 3.61±1.19, after: 2.83±0.92; t(17) = 2.96, p = 0.009, dz = 0.718), but remained stable after wakefulness (mean±SD, before: 3.17±1.25, after: 3.33±1.14; t(17) = 0.55, p = 0.59, dz = 0.132), which resulted in a significant Intervention x PrePost interaction (F(1,17) = 5.36, p = 0.033, η^2^ = .24). However, post-hoc analyses revealed that subjective sleepiness did not differ significantly between sleep and wake days before the intervention (t(17) = 1.29, p = 0.215, dz = 0.312) as well as after the intervention (t(17) = 1.84, p = 0.083, dz = 0.447).

A different pattern was observed in the psychomotor vigilance test. The reaction times increased slightly after wakefulness (mean±SD, before: 355±14, after: 366±18 ms; t(17) = 1.89, p = 0.075, dz = 0.46) but remained stable after the sleep interval (mean±SD, before: 357±19, after: 352±15 ms; t(17) = 0.756, p = 0.46, dz = 0.183), leading to a significant Intervention x PrePost interaction (F(1,17) = 5.65, p = 0.029, η^2^ = .249). Post-hoc tests revealed comparable vigilance performance between sleep and wake days before the intervention (t(17) = 0.38, p = 0.71, dz = 0.093), but faster reaction times on sleep days after the intervention (t(17) = 2.83, p = 0.012, dz = 0.686).

### Event-related potentials

#### Differential conditioned responses in ERPs

In the first step of the ERP analysis, we took the average of the CS+ and CS-waveforms and compared them with ERPs to Standards at three channels typically used in ERP analyses: Fz, Cz, and Pz. We ran a manipulation check to test for the expected oddball effect (i.e., CSs as rare deviant vs frequent standard stimuli). This comparison between standard and deviant responses was done to check the validity of our design. As expected, a strong oddball effect was present in the ERP components N1, P3a, and N3 (Figure 2a; Tables S2-S6: p < 0.01 for main effects of Stimulus (collapsed CSs vs. standards) with large effect sizes and central accentuation (Stimulus x Channel interaction). P3b was significantly larger to CSs only at Pz, and the amplitude of P2 was slightly smaller to CSs, with the latter finding possibly reflecting a mismatch negativity to the CS deviants. Thus, the amplitude of all ERP components of interest was more pronounced (except P2) in response to rare stimuli serving as CSs than to the frequent standard stimuli (for a detailed description of the analyses see Supplementary materials).

**Figure 2.**
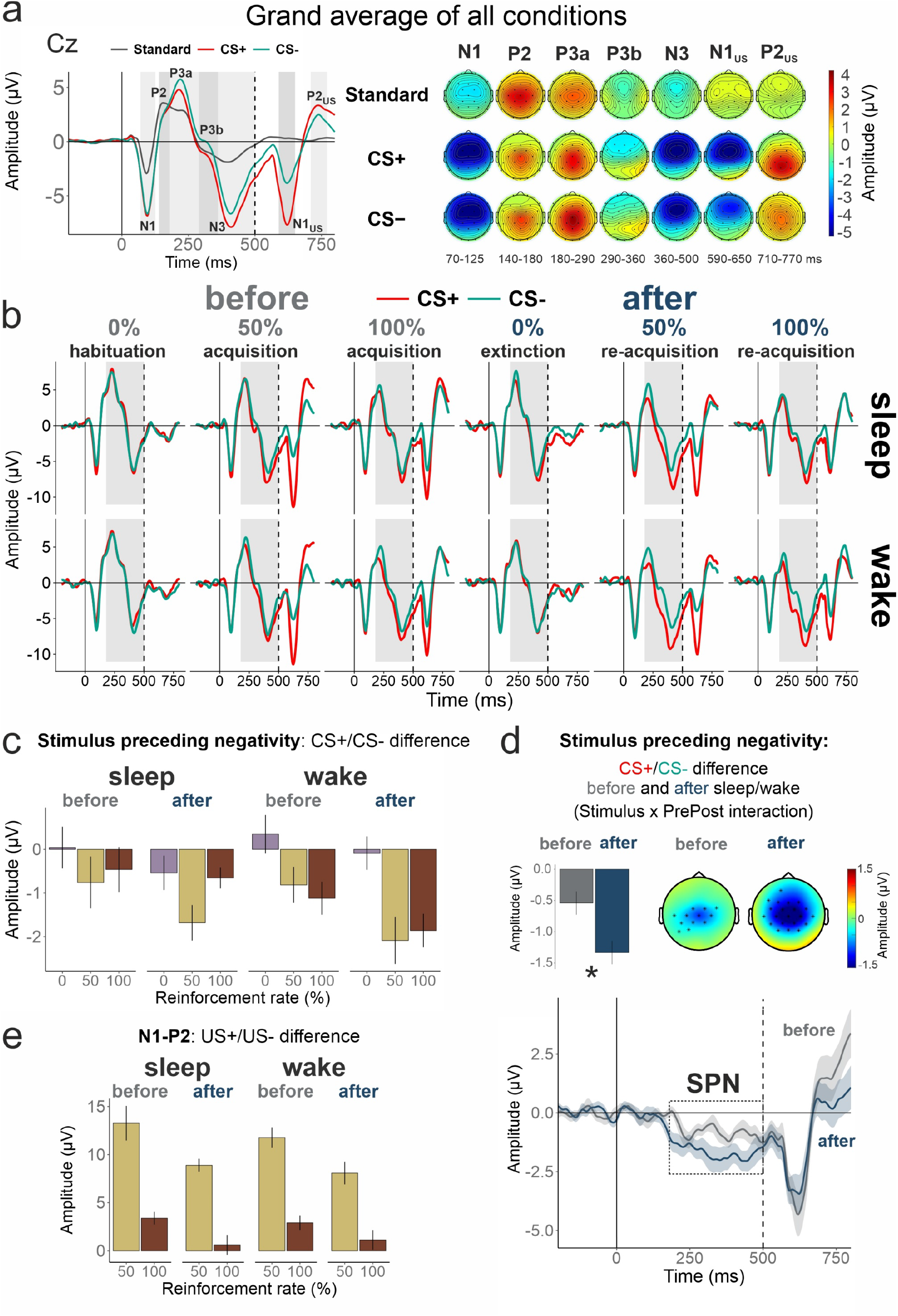
(a) Overall responses to the rare CS+ and CS-compared to frequent standard stimuli. Left panel: ERP curves at Cz channel. The grey boxes indicate (from left to right): N1, P2, P3a, P3b, N3, N1_US_, and P2_US_ time windows. Right panel: topographical maps with averaged ERPs in the corresponding time windows. (b) ERP curves at Cz channel for CS+ and CS-separately for the sleep and wake condition as well as before and after the intervention and for the different reinforcement rates. The grey box indicates the stimulus preceding negativity (SPN) time window (180-500 ms). (c) Barplots of the CS+/CS-difference in the SPN time window in all conditions. (d) CS+/CS-difference before and after the intervention (time effect). Top panels: average SPN in Cz, C3, and C4 (bar plot), and topoplots with black dots indicating significant clusters from the cluster-based permutation tests over all channels averaged in the SPN time window. Bottom panel: CS+/CS-difference in time. The shading indicates the standard error of the mean. (e) Barplots of the US+/US-difference in P2-N1 amplitude. The error bars represent the standard errors of the mean.

In the second step, we directly compared the CS+ and CS-ERP waveforms in the same time windows. The initial analysis revealed a significant Stimulus effect in P3a, P3b, and N3 time windows. A visual inspection revealed, however, that the effects were not separate for distinct time windows, but rather that there was one slow negative potential shift starting at about 180 ms after conditioned stimulus onset overlapping all faster components. The difference between CS+ and CS-amplitude significantly correlated between P3a, P3b, and N3 components (P3a-P3b: r = 0.7, p = 0.001; P3a-N3: r = 0.75, p < 0.001; P3b-N3: r = 0.87, p < 0.001) but not with N1 and P2. This slow shift is reminiscent of the stimulus preceding negativity (SPN (Böcker et al., 1994, 2001)). Therefore, we focused on the average SPN amplitude between 180 and 500 ms post-stimulus in the main analyses exploring the effects of sleep/wake on fear extinction and re-acquisition.

Before further statistical analyses, we assessed the spatial distribution of the Stimulus effect (CS+ vs CS-) via cluster-based permutation tests at all channels and time points. The tests revealed a stable presence of the Stimulus effect at central channels (see Figure S2 in the supplementary materials) over about 180-650 ms time interval. Confirming our assumption concerning the nature of this ERP component, this distribution was more typical for the SPN than for any of the faster ERP components in an oddball paradigm (Böcker et al., 1994; Kotani et al., 2015). Thus, the region of interest (ROI) for SPN analyses included C3, C4, and Cz channels (note that exploratory analyses using other channels around vertex yielded similar results).

Then we directly examined responses to the fear-conditioned stimulus (i.e. CS+ vs. CS-) during acquisition and re-acquisition (i.e., blocks 2 and 3 before the sleep/wake period and blocks 5 and 6 after the sleep/wake period). With regard to our first hypothesis, we observed a reliable correlate of differential fear conditioning in a slow wave spreading over a time window of 180 to at least 500 ms (F(1,17) = 20.52, p < 0.001, η^2^ = 0.547 for the main effect of Stimulus; Table S7), reflecting the stimulus-preceding negativity (SPN) (Figure 2b).

#### Effects of time and sleep on stimulus preceding negativity

In line with our second hypothesis, in a comparison of conditioned stimuli during acquisition and re-acquisition phases before and after the interventions, there was a pronounced time effect on the SPN, with the difference between the CS+ and CS-becoming larger after the retention interval (F(1,17) = 6.59, p = 0.02, η^2^ = 0.279 for Stimulus x PrePost interaction; Figures 2c,d, Table S7). Contrary to our third hypothesis, this effect was not differentially affected by whether subjects had a period of sleep or wakefulness during the interval (F(1,17) = 0.93, p = 0.348, η^2^ = 0.052 for Intervention x Stimulus x PrePost interaction). Generally, the amplitude of the SPN was larger after than before the intervention, larger in the wake than in the sleep condition, and larger with 100% reinforcement than with 50% reinforcement (F(1,17) = 16.23, 4.46, and 4.79, p < 0.001, 0.05, and 0.043, η^2^ = 0.488, 0.208, and 0.22 for main effects of PrePost, Intervention, and Block, respectively).

We then examined the effects of time and sleep on extinction learning during the first block after the sleep/wake period (block 4) compared to habituation before the sleep/wake interval (block 1). We did not observe a time or sleep effect in fear extinction. Although the SPN to both CSs was overall more negative during extinction than during habituation (F(1,17) = 52.81, p < 0.001, η^2^ = 0.756 for main effect PrePost), there was no difference between the CS+ and CS-nor between sleep and wakefulness or time points (p > 0.30 for all remaining main effects and interactions of Stimulus, PrePost and Intervention; Figure 2c, Table S8). To examine whether the lacking effects of sleep and time resulted from the large number of CS presentations (60 trials), we conducted an analysis of only the first 10 trials of extinction learning. However, the results remained essentially the same (Table S9), meaning that by the tenth trial the extinction has already happened.

#### Unconditioned responses

To test whether sleep or time might affect the processing of unconditioned stimuli, we compared the responses between US+ and US-in the acquisition and re-acquisition blocks. Overall, the US+ elicited a larger amplitude of the N1-P2 difference as compared to US- (F(1,17) = 35.98, p < 0.001, η^2^ = 0.679; Figure 2e; Table S10). Both unconditioned responses in general and the US+/US-difference decreased over time from before to after the sleep/wake interval (F(1,17) = 21.21, p < 0.001, η^2^ = 0.555 for main effect of PrePost, F(1,17) = 11.84, p = 0.003, η^2^ = 0.41 for PrePost x Stimulus interaction), speaking for a habituation effect independent of sleep (p > 0.20 for all effects of Intervention). Furthermore, unconditioned responses in general and the US+/US-difference were stronger in 50% compared to 100% reinforcement blocks, leading to a significant main effect of Block (F(1,17) = 92.02, p < 0.001, η^2^ = 0.844) and a significant Stimulus x Block interaction (F(1,17) = 64.89, p < 0.001, η^2^ = 0.792; Figure 2e).

### Subjective arousal and valence ratings

As expected, the valence ratings for the CS+ and CS-did not significantly differ after the habituation block (Figure 3; t(17) = 0.529, p = 0.60, dz = 0.128). Successful fear conditioning shifted ratings toward more negative valence for CS+ (F(1,17) = 4.61, p = 0.037, η^2^ = 0.213 for Block x Stimulus interaction and F(1,17) = 14.87, p = 0.001, η^2^ = 0.467 for main effect of Stimulus; Table S11) with no difference between 50% and 100% reinforcement rate (Stimulus effect after 50%: t(17) = 3.95, p = 0.001, dz = 0.958 and after 100% reinforcement rate blocks: t(17) = 3.37, p = 0.004, dz = 0.817; Table S12). Moreover, valence for CS-remained on a similar level over duration of the experiment (no significant effect of Block: F(1.3, 22.10) = 1.72, p = 0.205, η^2^ = 0.092). The arousal ratings were similar in dynamics than the valence ratings but not significant (Tables S13 and S14). Neither valence nor arousal ratings were affected by time or intervening sleep (all p > 0.20 for main effects and interactions with the factors PrePost and Intervention). However, exploratory correlation analysis revealed that more negative differential valence ratings of the CS+ during re-acquisition were associated with higher amounts of N2 sleep in the sleep condition (r = −0.69, p = 0.001; Table S1).

**Figure 3.**
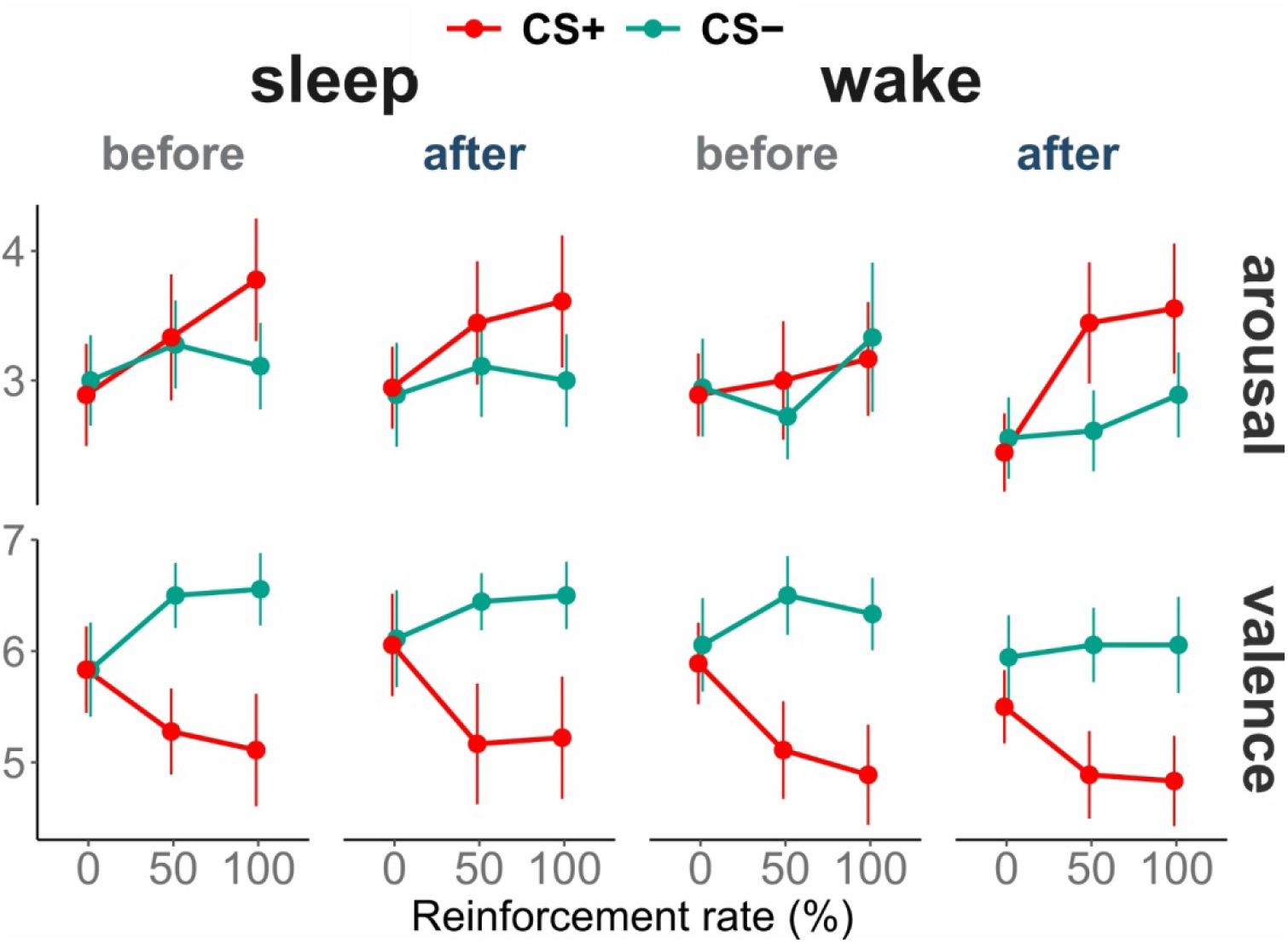
Valence and arousal ratings to CS+ and CS-collected after each block.

An analysis of responses to the US revealed more negative valence and higher arousal in US+ than US-ratings (F(1,17) = 28.87 and 122.92, p < 0.001 and < 0.001, η^2^ = 0.629 and 0.878 for main effect of Stimulus on arousal and valence, respectively; Tables S15 and S16). Additionally, arousal ratings decreased over time for both USs (F(1,17) = 6.94, p = 0.017, η^2^ = 0.29 for main effect of PrePost; mean±SD, before_arousal_: 5.15±0.99, after_arousal_: 4.71±1.27). This effect was not evident for valence ratings (p = 0.899; mean±SD, befor_evalence_: 4.80±1.06, after_valence_: 4.79±1.09).

Both ratings of the USs were not affected by intervening sleep (p > 0.05 for all effects of Intervention).

## Discussion

In our study, we investigated the effects of time and sleep on behavioral and central neural indices of fear learning in humans. The data showed a strong modulation of ERPs during fear learning, with a larger stimulus preceding negativity (SPN) to the CS+ compared to the CS-, indicating successful conditioning. Thus, the first hypothesis of our study was confirmed, highlighting the SPN as a viable neural (EEG) correlate of fear conditioning. Paralleling the neural findings, subjective ratings indicated more negative valence for the previously neutral CS+ after having been paired with a loud noise. After extinction training, valence ratings returned to the baseline level, and similarly the difference between CS+ and CS- in the SPN vanished after very few extinction trials. Importantly, after the retention interval (2 hours of sleep or wake), re-acquisition led to stronger neural conditioned responses compared to the original acquisition, evidenced by a larger SPN difference between the CS+ and CS-during reacquisition. This finding can be interpreted as confirmation of our second hypothesis (i.e., a time effect). However, both neural and behavioral conditioned responses were not affected by sleep, neither for fear extinction nor for fear re-acquisition. Our third hypothesis, therefore, was not supported by the data.

The finding that fear-conditioned neural evoked responses were reflected in a larger SPN probably indicates expectation and possible preparatory processes similar to flight or fight responses. An analogous increased SPN waveform has been shown to be predictive of emotionally charged events (Baas et al., 2002; Böcker et al., 2001; Dahl et al., 2020; Ferreira de Sá et al., 2019; Hellwig et al., 2008; Regan & Howard, 1995), while anticipatory attention to stimuli with no motivational value elicited reduced SPN (Brunia et al., 2011). This interpretation is in line with the observation that in the present study the SPN amplitude to the CS+ was generally larger in 100% reinforcement trials than in 50% reinforcement trials, whereas the responses to the US were weaker in 100% than 50% reinforcement trials. Possibly, the larger SPN in 100% reinforcement trials indicated stronger preparatory processes that, in turn, resulted in less pronounced unconditioned responses (MacNamara & Barley, 2018; Seidel et al., 2015).

Interestingly, our data did not reveal any effect of fear conditioning on early sensory ERP components such as N1 or P2. Instead, fear conditioning in our paradigm appeared to involve higher-order cognitive processing related to the anticipation of the unpleasant event. The absence of sensory or perceptual effects in conditioning contradicts some previous studies (Miskovic & Keil, 2012). Considering a pretty large number of averages in our paradigm (about 480 per condition per participant), the absence of these effects is unlikely to be caused by insufficient signal-to-noise ratio. Moreover, earlier studies argued that long-term (over days) and many trial paradigms are likely to affect early ERP components down to C1 in visual and P1 in auditory paradigms (Bublatzky & Schupp, 2012; Kluge et al., 2011; Stolarova et al., 2006). A possible factor contributing to this discrepancy may be an effect of latent inhibition, suggesting that unreinforced exposure of the to-be-conditioned stimulus slows down further associative learning with the same stimulus (Lubow, 1973). Our paradigm includes a relatively long habituation period preceding the acquisition block, which could have facilitated the engagement of higher order cognitive processes to overcome the effect of latent inhibition and to foster acquisition. In keeping with this explanation, other EEG studies that employed a habituation block immediately followed by the acquisition phase showed only late ERP effects (Bacigalupo & Luck, 2018; Danon-Kraun et al., 2021; Panitz et al., 2015; Stolz et al., 2019) as opposed to studies without the habituation phase (Bublatzky & Schupp, 2012; Pavlov & Kotchoubey, 2019; Sperl et al., 2021).

The enhanced SPN effect during re-acquisition (i.e., a larger SPN difference between the CS+ and CS-during re-acquisition compared to acquisition) can be interpreted as an indication of the consolidation of fear memories during the retention interval (i.e., over time, hypothesis 2). Notably, the SPN effect during acquisition before the intervention reached a plateau after the second block (with 50% reinforcement) and did not increase anymore in the 3rd block (with 100% reinforcement), suggesting that the initial learning process was already completed in the 2nd block. Therefore, the increase of the effect after intervention can most likely be interpreted as a result of (offline) consolidation processes rather than a simple continuation of learning. Furthermore, the effect was absent in the first block after intervention (no reinforcement), indicating successful extinction. Without consolidation, one would expect a completely new learning process after the retention interval that should be similar to the original fear memory acquisition. The finding that re-acquisition after the interval was more pronounced than original fear memory acquisition, suggests that offline consolidation strengthened the underlying memory traces which resulted in a facilitation of re-acquisition. However, we cannot fully rule out that alternative processes such as latent continuation of learning may have affected our results at least partially.

The finding that fear memory consolidation over time was only evident in neural responses in the present study, but not in subjective ratings of valence and arousal, suggests that neural responses are more sensitive to fear memory consolidation effects than behavioral measures. Thus, future studies should include such neural measures like ERP to capture subtle effects of consolidation that may be overlooked by less sensitive behavioral measures.

As stated in the introduction, the evidence of sleep benefits for emotional memory consolidation is mixed, and the results are controversial (e.g., Schäfer et al., 2020; Lerner et al., 2021), even those obtained by the same group (e.g., Menz et al., 2013; Menz et al., 2016). Nevertheless, basing on the findings that sleep consistently strengthens consolidation of other kinds of memory (Diekelmann & Born, 2010; Schimke et al., 2021; Schmid et al., 2020), we hypothesized that such positive effects would also be found on the consolidation of fear memory as well. This hypothesis was not confirmed. Our findings did not reveal any signs of differential fear conditioning after a short nap sleep compared to wakefulness. This is, however, in line with some previous studies that also did not observe stronger fear responses after sleep compared to wakefulness, e.g., in skin conductance responses (Zenses et al., 2020). Animal research may provide some insight into the mixed findings, suggesting that sleep plays a greater role in the consolidation of tasks that depend on the hippocampus, such as context conditioning, whereas tasks that mainly rely on the amygdala, such as cue conditioning, benefit to a lesser extent from sleep (Graves, 2003; Hagewoud et al., 2010; Phillips & LeDoux, 1992). Considering that the present paradigm is more akin to cue conditioning (i.e., a tone-cue preceding the US), the lack of a sleep effect might be explained by a relatively weak involvement of the hippocampus. Alternatively, the lacking effect of sleep on extinction learning might have been due to too many extinction trials in the present study (60 trials). However, additional analyses of only the first 10 extinction trials likewise did not reveal any differences between the sleep and wake condition. Yet another possibility is that sleep effects may be evident only in later responses, i.e., after 500 ms. However, again, additional analyses also showed no significant differences in time windows after 500 ms.

Overall, in the current study, we examined the neural mechanisms underlying the formation and consolidation of fear memories, with a particular focus on changes in fear responses over time and sleep. Applying sensitive electrophysiological measures, we provide evidence that fear memories are consolidated over time, with this effect being independent of sleep. These findings suggest that pathological fear responses as a result of exposure to strong fear-evoking events may increase over time, possibly contributing to the development of anxiety-related disorders. Sleep, however, does not seem to influence this process, neither enhancing nor reducing fear responses. This new understanding of the evolution of fear memories over time should be taken into account in the prevention and treatment of anxiety-related disorders such as PTSD and specific phobias.

## Limitations

Several limitations of the present study should be considered. First, our sample size was too small to permit an in-depth regression or correlation analysis to examine the possible effects of particular sleep stages (e.g., REM) on subsequent fear extinction and re-acquisition. The role of certain sleep stages for the consolidation of fear conditioning is an important question, but much larger samples would be needed to run feasible analyses. Second, we applied a nap design, which has the advantage that sleep and wake groups were comparable with regard to circadian timing. However, while a number of studies found that even short naps can facilitate memory consolidation like night sleep (e.g., Lahl et al., 2008; Sopp et al., 2018), other studies found no benefits of naps (McDevitt et al., 2018; Zenses et al., 2020). Thus, longer sleep periods and particularly a full night of sleep may have different effects on fear memory consolidation. Third, we did not assess expectancy ratings, i.e., how strongly participants expect the US to follow the CS+, which have frequently been employed in human fear conditioning research, and might be informative with regard to associations of these measures with neural fear responses. Forth, neural measures as conditioned responses possess great advantages as they allow detecting the unfolding of brain processes in real time with a perfect time resolution. However, the corresponding disadvantage is the necessity to average a large number of trials, resulting in the impossibility to examine responses in single trials. Finally, a serious limitation is the lack of peripheral physiological measures such as skin conductance and heart rate. This, however, was determined by the experimental design, as using peripheral physiological signals within the same design would have resulted in much longer durations of the experimental sessions due to the slower development of peripheral responses. Considering that the experimental sessions were already quite straining for our subjects, a further prolongation would have led to severe problems with monotony and fatigue. At the same time, our data open the way to overcome this limitation in future studies: because the SPN in our study demonstrated a reasonably high signal-to-noise ratio, future studies may employ concurrent EEG and peripheral physiology recordings by reducing the number of blocks and the number of trials per block.

## Acknowledgements

The study was supported by the German Research Society (Deutsche Forschungsgemeinschaft, DFG), grant KO-1753/13-4. The authors thank Martin King, Marina Zimmermann, and Nina Heidemann for their help with data collection.

## Author contributions statement

Yuri G. Pavlov: Conceptualization, Investigation, Data curation, Formal analysis, Project administration, Visualization, Supervision, Methodology, Writing - original draft, Writing - review & editing.

Nadezhda V. Pavlova: Conceptualization, Investigation, Formal analysis, Writing - review & editing.

Boris Kotchoubey: Conceptualization, Funding acquisition, Project administration, Methodology, Writing - review & editing.

Susanne Diekelmann: Conceptualization, Methodology, Writing - review & editing.

## Data availability statement

The data are publicly available on OSF (https://osf.io/xph69/).

## Footnotes

1 Please note that using all instead of these a-priori selected three channels would result in the same time window selection

