## Supplementary for "Fear memory in humans is consolidated over time independent of sleep"

### Supplementary methods

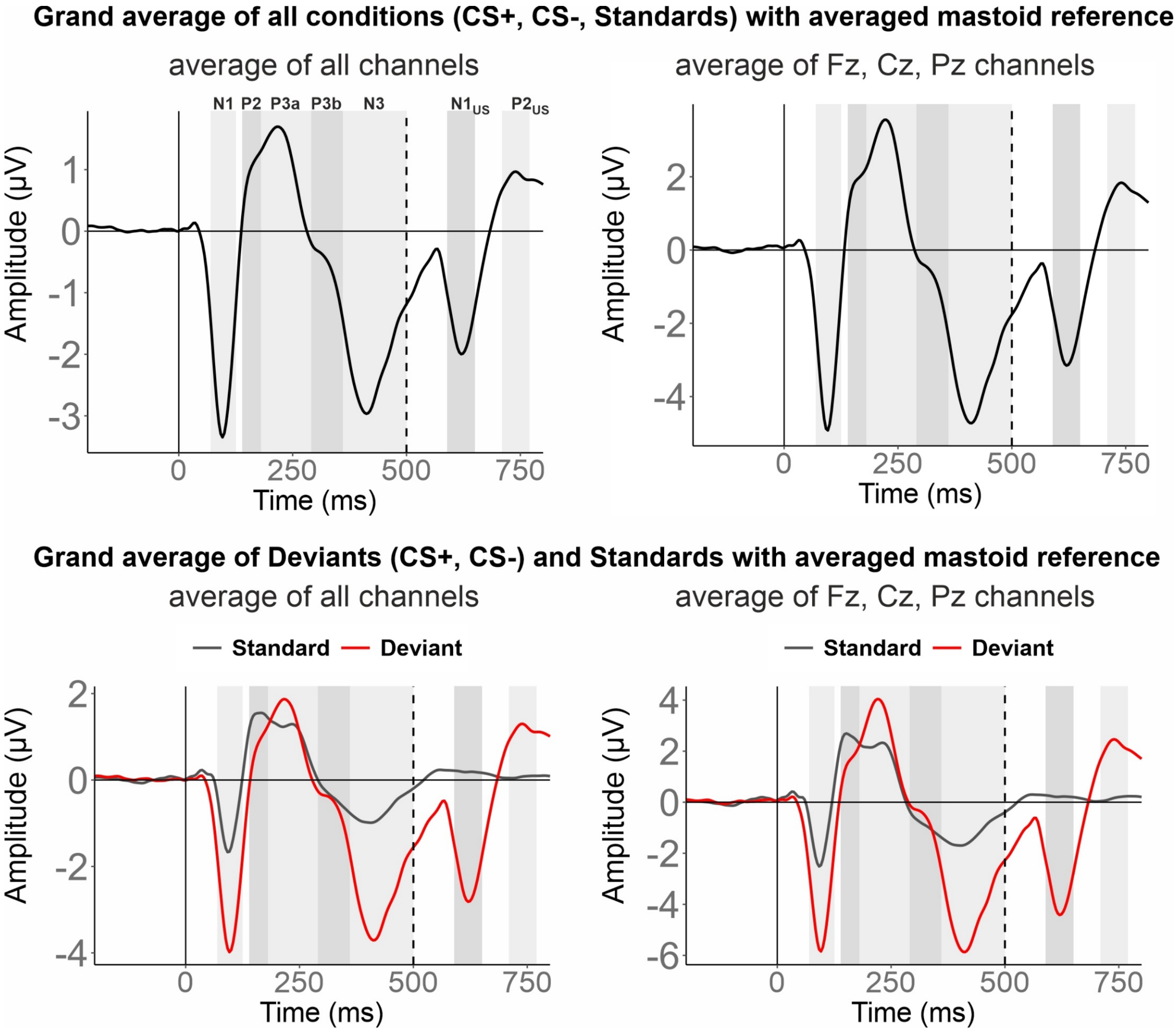


Figure S1 – The flatten average of all conditions (upper panels) or collapsed CSs and Standards separately (bottom panels) using all channels (left panels) or Fz, Cz, Pz channels (right panels). The upper right figure was used for selection of the time windows for the first analysis comparing Standard and Deviant stimuli (see Oddball effect section below). Time windows are marked with grey bars (left to right: N1, P2, P3a, P3b, N3, $N1_{US}$, $P2_{US}$ time windows).

### Supplementary results

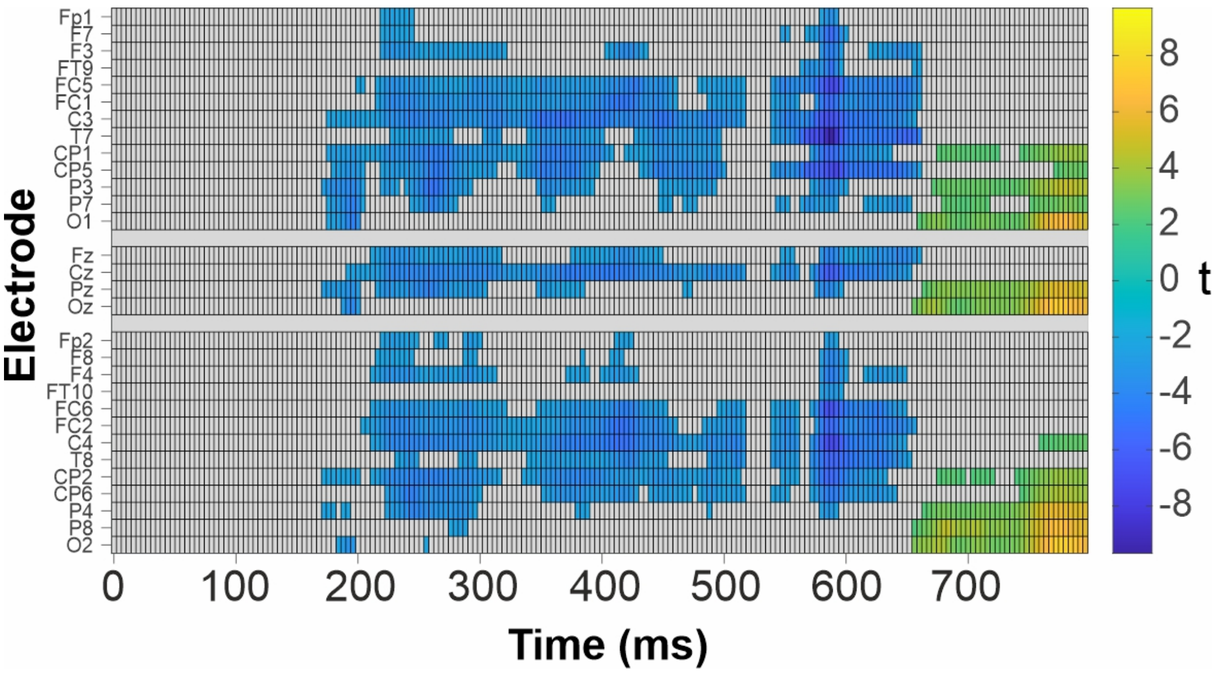


Figure S2 – Raster plot illustrating the results of the cluster-based permutation tests of CS+ vs CS- comparison at all channels and time points in 50% and 100% reinforcement rate blocks combined.

Table S1

*Correlation between conditioning indices (CS+/CS- difference) and time spent in the different sleep stages*

| Variable | *M* | *SD* | 1 | 2 | 3 | 4 | 5 | 6 | 7 | 8 |
|---|---|---|---|---|---|---|---|---|---|---|
| 1. N2, min | 43.00 | 17.36 | | | | | | | | |
| 2. N3, min | 19.89 | 15.96 | -0.01 | | | | | | | |
| 3. REM, min | 9.94 | 11.54 | **0.6*** | -0.18 | | | | | | |
| 4. SPN, µV extinction | -0.54 | 1.69 | 0.37 | -0.3 | 0.42 | | | | | |
| 5. SPN, µV re-acquisition | -1.17 | 1.52 | 0.03 | -0.17 | 0.28 | 0.34 | | | | |
| 6. Arousal extinction | 0.06 | 1.11 | -0.23 | -0.15 | -0.32 | -0.32 | 0.29 | | | |
| 7. Valence extinction | -0.06 | 2.34 | -0.38 | 0 | -0.21 | 0.02 | -0.06 | 0.11 | | |
| 8. Arousal re-acquisition | 0.47 | 1.91 | -0.14 | -0.12 | -0.24 | -0.26 | 0.22 | **0.73*** | 0.16 | |
| 9. Valence re-acquisition | -1.28 | 2.65 | **-0.69*** | -0.06 | -0.24 | -0.01 | -0.12 | -0.17 | **0.54*** | -0.07 |

*Note. M* and *SD* are used to represent mean and standard deviation, respectively. * indicates $p < .05$.

#### Oddball effect

*N1, P2, P3a, P3b, N3 time windows. ANOVA with factors: Intervention: Sleep/Wake; PrePost: Before/After; Stimulus: Standard/Deviant; Block: 0%/50%/100% reinforcement rate blocks; Channel: Fz/Cz/Pz.*

In these supplementary analyses, since the analysis involved a great number of comparisons (31 possible effects by 5 components), all effects, which were not strongly expected (i.e., explorative), were regarded as significant only when $p < 0.01$. N1 and N3 significantly increased from Block1 to Block3, while positive components decreased. The P3a decrease was particularly strong at Cz (Block x Channel interaction). As the P3b amplitude to standards was virtually zero, the decrease manifested itself only in the response to deviants at Pz, yielding highly significant interactions between Block and Stimulus, Block and Channel, and between Block, Stimulus and Channel.

The positive components generally decreased after the intervention (main effect of PrePost). Significant PrePost x Stimulus interactions indicated that after the sleep/wake period, the standard/deviant difference for P3a (i.e., more positive amplitude to deviant than standard) decreased, but the standard/deviant differences for P2 and N3 (with more positive amplitudes to standard than deviant) increased. In simpler words, this means that time had a stronger effect on deviant responses than on standard responses. Finally, the standard/deviant difference in P2 appeared to be larger in the wake condition than in the sleep condition; however, because it was a singular Intervention effect, and because this was only a Stimulus x Intervention interaction and not (as might have been expected) a Stimulus x PrePost x Intervention interaction, the finding should be considered with caution.

Table S2 – ANOVA of N1 amplitude

| Predictor | $df_{Num}$ | $df_{Den}$ | $F$ | $p$ | $\eta^2$ |
|---|---|---|---|---|---|
| Intervention | 1 | 17 | 0.02 | 0.888 | 0.001 |
| PrePost | 1 | 17 | 2.42 | 0.138 | 0.125 |
| Stimulus | 1 | 17 | 143.13 | **<.001** | 0.894 |
| Block | 1.54 | 26.22 | 16.78 | **<.001** | 0.497 |
| Channel | 1.1 | 18.71 | 11.37 | **0.003** | 0.401 |
| Intervention x PrePost | 1 | 17 | 0.01 | 0.93 | <.001 |
| Intervention x Stimulus | 1 | 17 | 0.4 | 0.534 | 0.023 |
| PrePost x Stimulus | 1 | 17 | 5.27 | **0.035** | 0.237 |
| Intervention x Block | 1.91 | 32.44 | 0.16 | 0.841 | 0.009 |
| PrePost x Block | 1.86 | 31.68 | 3.85 | **0.035** | 0.184 |
| Stimulus x Block | 1.7 | 28.87 | 1.27 | 0.292 | 0.069 |
| Intervention x Channel | 1.42 | 24.15 | 0.78 | 0.429 | 0.044 |
| PrePost x Channel | 1.4 | 23.83 | 0.23 | 0.718 | 0.013 |
| Stimulus x Channel | 1.2 | 20.37 | 69.88 | **<.001** | 0.804 |
| Block x Channel | 1.97 | 33.54 | 2.39 | 0.108 | 0.123 |
| Intervention x PrePost x Stimulus | 1 | 17 | 0.1 | 0.759 | 0.006 |
| Intervention x PrePost x Block | 1.96 | 33.33 | 0.59 | 0.557 | 0.034 |
| Intervention x Stimulus x Block | 1.85 | 31.39 | 0.85 | 0.427 | 0.048 |
| PrePost x Stimulus x Block | 1.84 | 31.21 | 1.41 | 0.259 | 0.076 |
| Intervention x PrePost x Channel | 1.28 | 21.76 | 0.04 | 0.894 | 0.002 |
| Intervention x Stimulus x Channel | 1.53 | 25.95 | 1.31 | 0.279 | 0.072 |

| | | | | | |
|---|---|---|---|---|---|
| PrePost x Stimulus x Channel | 1.24 | 21.1 | 0.95 | 0.362 | 0.053 |
| Intervention x Block x Channel | 2.3 | 39.17 | 0.92 | 0.417 | 0.052 |
| PrePost x Block x Channel | 2.17 | 36.83 | 3.61 | **0.034** | 0.175 |
| Stimulus x Block x Channel | 1.93 | 32.86 | 2.17 | 0.131 | 0.113 |
| Intervention x PrePost x Stimulus x Block | 1.62 | 27.54 | 0.9 | 0.398 | 0.05 |
| Intervention x PrePost x Stimulus x Channel | 1.38 | 23.52 | 0.86 | 0.398 | 0.048 |
| Intervention x PrePost x Block x Channel | 1.85 | 31.42 | 0.16 | 0.835 | 0.009 |
| Intervention x Stimulus x Block x Channel | 2.31 | 39.24 | 0.41 | 0.696 | 0.024 |
| PrePost x Stimulus x Block x Channel | 2.3 | 39.17 | 1 | 0.385 | 0.056 |
| Intervention x PrePost x Stimulus x Block x Channel | 2.48 | 42.14 | 1.31 | 0.282 | 0.072 |

Table S3 – ANOVA of P2 amplitude

| Predictor | $df_{Num}$ | $df_{Den}$ | $F$ | $p$ | $\eta^2$ |
|---|---|---|---|---|---|
| Intervention | 1 | 17 | 1.37 | 0.258 | 0.075 |
| PrePost | 1 | 17 | 10.45 | **0.005** | 0.381 |
| Stimulus | 1 | 17 | 9.74 | **0.006** | 0.364 |
| Block | 1.51 | 25.74 | 14.63 | **<.001** | 0.462 |
| Channel | 1.82 | 30.96 | 18.81 | **<.001** | 0.525 |
| Intervention x PrePost | 1 | 17 | 0.16 | 0.692 | 0.009 |
| Intervention x Stimulus | 1 | 17 | 8.56 | **0.009** | 0.335 |
| PrePost x Stimulus | 1 | 17 | 9.26 | **0.007** | 0.353 |
| Intervention x Block | 1.41 | 23.93 | 1.38 | 0.264 | 0.075 |
| PrePost x Block | 1.5 | 25.5 | 1.27 | 0.29 | 0.069 |
| Stimulus x Block | 1.39 | 23.68 | 0.21 | 0.733 | 0.012 |
| Intervention x Channel | 1.28 | 21.77 | 0.15 | 0.765 | 0.009 |
| PrePost x Channel | 1.24 | 21.06 | 6.32 | **0.015** | 0.271 |
| Stimulus x Channel | 1.58 | 26.81 | 4.84 | **0.022** | 0.221 |
| Block x Channel | 2.02 | 34.34 | 3.70 | **0.035** | 0.179 |
| Intervention x PrePost x Stimulus | 1 | 17 | 0.74 | 0.403 | 0.041 |
| Intervention x PrePost x Block | 1.64 | 27.83 | 0.05 | 0.923 | 0.003 |
| Intervention x Stimulus x Block | 1.87 | 31.81 | 0.1 | 0.895 | 0.006 |
| PrePost x Stimulus x Block | 1.88 | 31.91 | 1.49 | 0.241 | 0.081 |
| Intervention x PrePost x Channel | 1.39 | 23.64 | 1.74 | 0.201 | 0.093 |
| Intervention x Stimulus x Channel | 1.58 | 26.87 | 1.12 | 0.328 | 0.062 |
| PrePost x Stimulus x Channel | 1.2 | 20.44 | 0.21 | 0.695 | 0.012 |
| Intervention x Block x Channel | 2.96 | 50.28 | 1.16 | 0.333 | 0.064 |
| PrePost x Block x Channel | 2.29 | 38.94 | 2.26 | 0.112 | 0.117 |
| Stimulus x Block x Channel | 1.44 | 24.45 | 0.07 | 0.874 | 0.004 |
| Intervention x PrePost x Stimulus x Block | 1.23 | 20.91 | 0.38 | 0.586 | 0.022 |
| Intervention x PrePost x Stimulus x Channel | 1.38 | 23.42 | 3.66 | 0.056 | 0.177 |
| Intervention x PrePost x Block x Channel | 1.74 | 29.58 | 1.69 | 0.204 | 0.091 |
| Intervention x Stimulus x Block x Channel | 2.44 | 41.49 | 0.16 | 0.890 | 0.009 |
| PrePost x Stimulus x Block x Channel | 2.59 | 44.07 | 1.23 | 0.308 | 0.067 |
| Intervention x PrePost x Stimulus x Block x Channel | 2.41 | 40.89 | 0.78 | 0.487 | 0.044 |

Table S4 – ANOVA of P3a amplitude

| Predictor | $df_{Num}$ | $df_{Den}$ | $F$ | $p$ | $\eta^2$ |
|---|---|---|---|---|---|
| Intervention | 1 | 17 | 3.33 | 0.085 | 0.164 |
| PrePost | 1 | 17 | 27.20 | **<.001** | 0.615 |
| Stimulus | 1 | 17 | 13.40 | **0.002** | 0.441 |
| Block | 1.82 | 30.88 | 37.51 | **<.001** | 0.688 |
| Channel | 1.81 | 30.74 | 18.16 | **<.001** | 0.516 |
| Intervention x PrePost | 1 | 17 | 3.21 | 0.091 | 0.159 |
| Intervention x Stimulus | 1 | 17 | 5.05 | **0.038** | 0.229 |
| PrePost x Stimulus | 1 | 17 | 15.55 | **0.001** | 0.478 |
| Intervention x Block | 1.63 | 27.63 | 0.42 | 0.620 | 0.024 |
| PrePost x Block | 1.95 | 33.21 | 1.71 | 0.197 | 0.091 |
| Stimulus x Block | 1.88 | 31.92 | 4.74 | **0.017** | 0.218 |
| Intervention x Channel | 1.55 | 26.39 | 0.11 | 0.851 | 0.006 |
| PrePost x Channel | 1.4 | 23.79 | 2.62 | 0.109 | 0.133 |
| Stimulus x Channel | 1.5 | 25.46 | 12.65 | **<.001** | 0.427 |
| Block x Channel | 2.37 | 40.33 | 8.96 | **<.001** | 0.345 |
| Intervention x PrePost x Stimulus | 1 | 17 | 0.92 | 0.351 | 0.051 |
| Intervention x PrePost x Block | 1.81 | 30.69 | 1.09 | 0.342 | 0.06 |
| Intervention x Stimulus x Block | 1.88 | 31.91 | 0.32 | 0.716 | 0.018 |
| PrePost x Stimulus x Block | 1.96 | 33.4 | 0.82 | 0.447 | 0.046 |
| Intervention x PrePost x Channel | 1.21 | 20.5 | 2.47 | 0.127 | 0.127 |
| Intervention x Stimulus x Channel | 1.64 | 27.84 | 1.45 | 0.249 | 0.079 |
| PrePost x Stimulus x Channel | 1.23 | 20.98 | 0.92 | 0.368 | 0.052 |
| Intervention x Block x Channel | 3.33 | 56.55 | 0.13 | 0.952 | 0.008 |
| PrePost x Block x Channel | 2.42 | 41.19 | 0.36 | 0.738 | 0.021 |
| Stimulus x Block x Channel | 2.12 | 36.07 | 4.38 | **0.018** | 0.205 |
| Intervention x PrePost x Stimulus x Block | 1.96 | 33.24 | 0.66 | 0.522 | 0.037 |
| Intervention x PrePost x Stimulus x Channel | 1.55 | 26.39 | 1.69 | 0.207 | 0.091 |
| Intervention x PrePost x Block x Channel | 2.39 | 40.55 | 1.15 | 0.335 | 0.063 |
| Intervention x Stimulus x Block x Channel | 1.8 | 30.67 | 4.21 | **0.028** | 0.198 |
| PrePost x Stimulus x Block x Channel | 2.73 | 46.42 | 0.37 | 0.756 | 0.021 |
| Intervention x PrePost x Stimulus x Block x Channel | 2.85 | 48.41 | 3.38 | **0.028** | 0.166 |

Table S5 – ANOVA of P3b amplitude

| Predictor | $df_{Num}$ | $df_{Den}$ | $F$ | $p$ | $\eta^2$ |
|---|---|---|---|---|---|
| Intervention | 1 | 17 | 2.49 | 0.133 | 0.128 |
| PrePost | 1 | 17 | 23.12 | **<.001** | 0.576 |
| Stimulus | 1 | 17 | 1.72 | 0.207 | 0.092 |
| Block | 1.47 | 24.91 | 36.46 | **<.001** | 0.682 |
| Channel | 1.54 | 26.1 | 15.57 | **<.001** | 0.478 |
| Intervention x PrePost | 1 | 17 | 0.86 | 0.366 | 0.048 |
| Intervention x Stimulus | 1 | 17 | 2.66 | 0.121 | 0.135 |
| PrePost x Stimulus | 1 | 17 | 40.62 | **<.001** | 0.705 |
| Intervention x Block | 1.88 | 31.98 | 0.41 | 0.655 | 0.024 |
| PrePost x Block | 1.96 | 33.3 | 0.36 | 0.698 | 0.021 |
| Stimulus x Block | 1.91 | 32.46 | 26.63 | **<.001** | 0.61 |
| Intervention x Channel | 1.25 | 21.27 | 0.85 | 0.39 | 0.048 |

| | | | | | |
|---|---|---|---|---|---|
| PrePost x Channel | 1.59 | 27.01 | 0.81 | 0.431 | 0.045 |
| Stimulus x Channel | 1.65 | 28.05 | 15.73 | **<.001** | 0.481 |
| Block x Channel | 2.2 | 37.39 | 7.02 | **0.002** | 0.292 |
| Intervention x PrePost x Stimulus | 1 | 17 | 2.31 | 0.147 | 0.12 |
| Intervention x PrePost x Block | 1.88 | 32.03 | 0.14 | 0.862 | 0.008 |
| Intervention x Stimulus x Block | 1.97 | 33.5 | 0.06 | 0.94 | 0.004 |
| PrePost x Stimulus x Block | 1.91 | 32.43 | 1.81 | 0.18 | 0.096 |
| Intervention x PrePost x Channel | 1.21 | 20.52 | 2.09 | 0.161 | 0.11 |
| Intervention x Stimulus x Channel | 1.32 | 22.47 | 0.47 | 0.554 | 0.027 |
| PrePost x Stimulus x Channel | 1.7 | 28.83 | 2.55 | 0.103 | 0.131 |
| Intervention x Block x Channel | 2.48 | 42.24 | 0.11 | 0.93 | 0.006 |
| PrePost x Block x Channel | 2.39 | 40.57 | 0.29 | 0.788 | 0.017 |
| Stimulus x Block x Channel | 2.84 | 48.35 | 9.39 | **<.001** | 0.356 |
| Intervention x PrePost x Stimulus x Block | 1.74 | 29.54 | 0.08 | 0.898 | 0.005 |
| Intervention x PrePost x Stimulus x Channel | 1.2 | 20.43 | 1.12 | 0.315 | 0.062 |
| Intervention x PrePost x Block x Channel | 2.25 | 38.32 | 0.27 | 0.788 | 0.016 |
| Intervention x Stimulus x Block x Channel | 2.72 | 46.29 | 1.21 | 0.315 | 0.066 |
| PrePost x Stimulus x Block x Channel | 2.73 | 46.42 | 0.79 | 0.497 | 0.044 |
| Intervention x PrePost x Stimulus x Block x Channel | 2.29 | 38.89 | 2.80 | 0.067 | 0.141 |

Table S6 – ANOVA of N3 amplitude

| Predictor | $df_{Num}$ | $df_{Den}$ | $F$ | $p$ | $\eta^2$ |
|---|---|---|---|---|---|
| Intervention | 1 | 17 | 0.66 | 0.427 | 0.037 |
| PrePost | 1 | 17 | 4.41 | 0.051 | 0.206 |
| Stimulus | 1 | 17 | 74.37 | **<.001** | 0.814 |
| Block | 1.37 | 23.3 | 12.94 | **<.001** | 0.432 |
| Channel | 1.56 | 26.45 | 48.03 | **<.001** | 0.739 |
| Intervention x PrePost | 1 | 17 | 0.02 | 0.894 | 0.001 |
| Intervention x Stimulus | 1 | 17 | 1.32 | 0.267 | 0.072 |
| PrePost x Stimulus | 1 | 17 | 14.62 | **0.001** | 0.462 |
| Intervention x Block | 1.74 | 29.55 | 1.74 | 0.195 | 0.093 |
| PrePost x Block | 1.75 | 29.81 | 5.49 | **0.012** | 0.244 |
| Stimulus x Block | 1.73 | 29.37 | 6.93 | **0.005** | 0.29 |
| Intervention x Channel | 1.39 | 23.55 | 2 | 0.167 | 0.105 |
| PrePost x Channel | 1.6 | 27.25 | 1.59 | 0.223 | 0.086 |
| Stimulus x Channel | 1.61 | 27.37 | 37.29 | **<.001** | 0.687 |
| Block x Channel | 2.04 | 34.7 | 3.03 | 0.06 | 0.151 |
| Intervention x PrePost x Stimulus | 1 | 17 | 0.35 | 0.564 | 0.02 |
| Intervention x PrePost x Block | 1.9 | 32.33 | 0.15 | 0.851 | 0.009 |
| Intervention x Stimulus x Block | 1.78 | 30.19 | 0.78 | 0.455 | 0.044 |
| PrePost x Stimulus x Block | 1.96 | 33.33 | 0.43 | 0.651 | 0.025 |
| Intervention x PrePost x Channel | 1.33 | 22.53 | 0.14 | 0.781 | 0.008 |
| Intervention x Stimulus x Channel | 1.34 | 22.86 | 0.96 | 0.365 | 0.053 |
| PrePost x Stimulus x Channel | 1.42 | 24.1 | 0.49 | 0.555 | 0.028 |
| Intervention x Block x Channel | 2.03 | 34.59 | 0.73 | 0.489 | 0.041 |
| PrePost x Block x Channel | 2.61 | 44.44 | 2.26 | 0.102 | 0.117 |
| Stimulus x Block x Channel | 2.26 | 38.37 | 3.85 | **0.026** | 0.185 |
| Intervention x PrePost x Stimulus x Block | 1.8 | 30.63 | 1.72 | 0.198 | 0.092 |

| | | | | | |
|---|---|---|---|---|---|
| Intervention x PrePost x Stimulus x Channel | 1.41 | 23.91 | 0.01 | 0.966 | <.001 |
| Intervention x PrePost x Block x Channel | 1.82 | 30.87 | 0.43 | 0.634 | 0.025 |
| Intervention x Stimulus x Block x Channel | 2.87 | 48.82 | 2.92 | **0.045** | 0.147 |
| PrePost x Stimulus x Block x Channel | 2.73 | 46.37 | 2.28 | 0.097 | 0.118 |
| Intervention x PrePost x Stimulus x Block x Channel | 2.47 | 41.99 | 2.31 | 0.101 | 0.120 |

#### Conditioned stimulus: stimulus preceding negativity analyses

*ANOVA of SPN at C3, C4, and Cz channels to conditioned stimuli during 50% and 100% reinforcement rate blocks, i.e. acquisition and re-acquisition (Intervention: Sleep/Wake; PrePost: Before/After; Stimulus: CS+/CS-; Block: 50%/100%)*

Table S7 – ANOVA of SPN to conditioned stimuli: acquisition/re-acquisition

| Predictor | $df_{Num}$ | $df_{Den}$ | $F$ | $p$ | $\eta^2$ |
|---|---|---|---|---|---|
| Intervention | 1 | 17 | 4.46 | **0.050** | 0.208 |
| PrePost | 1 | 17 | 16.23 | **<.001** | 0.488 |
| Stimulus | 1 | 17 | 20.52 | **<.001** | 0.547 |
| Block | 1 | 17 | 4.79 | **0.043** | 0.220 |
| Intervention x PrePost | 1 | 17 | 1.41 | 0.252 | 0.076 |
| Intervention x Stimulus | 1 | 17 | 1.46 | 0.243 | 0.079 |
| PrePost x Stimulus | 1 | 17 | 6.59 | **0.020** | 0.279 |
| Intervention x Block | 1 | 17 | 1.13 | 0.302 | 0.062 |
| PrePost x Block | 1 | 17 | 1.61 | 0.221 | 0.087 |
| Stimulus x Block | 1 | 17 | 2.56 | 0.128 | 0.131 |
| Intervention x PrePost x Stimulus | 1 | 17 | 0.93 | 0.348 | 0.052 |
| Intervention x PrePost x Block | 1 | 17 | 0.11 | 0.746 | 0.006 |
| Intervention x Stimulus x Block | 1 | 17 | 1.94 | 0.181 | 0.103 |
| PrePost x Stimulus x Block | 1 | 17 | 1.00 | 0.331 | 0.056 |
| Intervention x PrePost x Stimulus x Block | 1 | 17 | 0.02 | 0.885 | 0.001 |

*ANOVA of SPN at C3, C4, and Cz channels to conditioned stimuli during 0% reinforcement rate blocks, i.e. habituation and extinction (Intervention: Sleep/Wake; PrePost: Before/After; Stimulus: CS+/CS-)*

Table S8 – ANOVA of SPN to conditioned stimuli: habituation/extinction (block1/block4)

| Predictor | $df_{Num}$ | $df_{Den}$ | $F$ | $p$ | $\eta^2$ |
|---|---|---|---|---|---|
| Intervention | 1 | 17 | 0.65 | 0.432 | 0.037 |
| PrePost | 1 | 17 | 52.81 | **<.001** | 0.756 |
| Stimulus | 1 | 17 | 0.16 | 0.699 | 0.009 |
| Intervention:PrePost | 1 | 17 | 0.16 | 0.696 | 0.009 |
| Intervention:Stimulus | 1 | 17 | 0.73 | 0.405 | 0.041 |
| PrePost:Stimulus | 1 | 17 | 1 | 0.330 | 0.056 |
| Intervention:PrePost:Stimulus | 1 | 17 | 0.08 | 0.778 | 0.005 |

Table S9 – ANOVA of SPN to conditioned stimuli: habituation/extinction, only first 10 trials

| Predictor | $df_{Num}$ | $df_{Den}$ | $F$ | $p$ | $\eta^2$ |
|---|---|---|---|---|---|
| Intervention | 1 | 17 | 0.88 | 0.362 | 0.049 |
| PrePost | 1 | 17 | 9.35 | **0.007** | 0.355 |
| Stimulus | 1 | 17 | 0.06 | 0.809 | 0.004 |
| Intervention:PrePost | 1 | 17 | 2.78 | 0.114 | 0.141 |
| Intervention:Stimulus | 1 | 17 | 0.16 | 0.692 | 0.009 |
| PrePost:Stimulus | 1 | 17 | 0.45 | 0.509 | 0.026 |
| Intervention:PrePost:Stimulus | 1 | 17 | 0.13 | 0.719 | 0.008 |

#### Later effects in unreinforced trials

To explore whether there were any effects of sleep or time beyond the analyzed 180-500 ms interval (e.g., late positive potential (LPP)), we selected only unreinforced trials from the 50% reinforcement rate block (because in reinforced trials the US was presented at 500 ms) and compared the CS+ and CS- over the whole duration of the trial (0-800 ms). As shown in Figure S2, there were no late ERPs that differentiated CS+ and CS- (cluster-based permutation tests did not reveal any significant clusters beyond 500 ms).

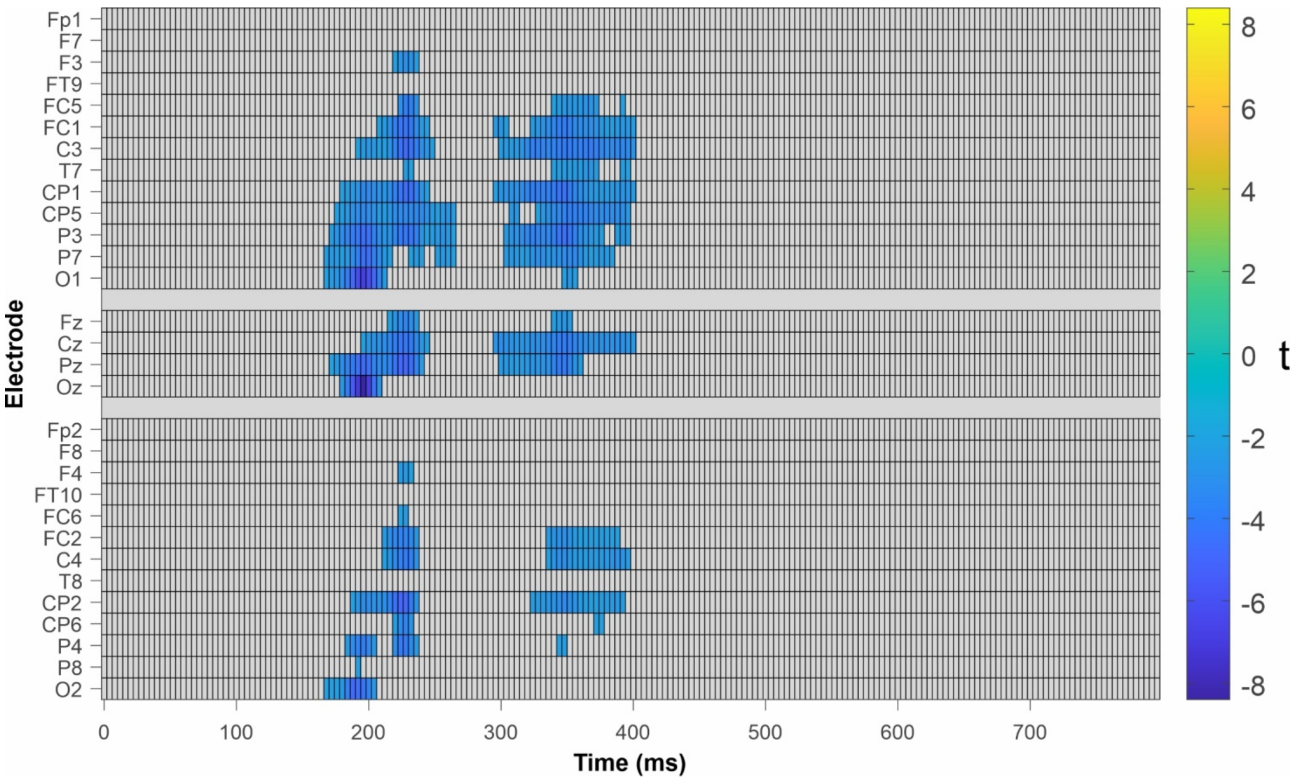


Figure S3 – Raster plot of the results of the cluster-based permutation tests CS+ vs CS- in 50% reinforcement rate block unreinforced trials only.

#### Unconditioned stimulus: N1-P2 difference analysis

*ANOVA with factors: Intervention: Sleep/Control; PrePost: Before/After; Stimulus: CS+/CS-; Block: 50% (only reinforced trials), 100% reinforcement rate blocks.*

Table S10 – ANOVA of N1-P2 amplitude difference at C3, C4, and Cz channels

| Predictor | $df_{Num}$ | $df_{Den}$ | $F$ | $p$ | $\eta^2$ |
|---|---|---|---|---|---|
| Intervention | 1 | 17 | 0.4 | 0.534 | 0.023 |
| PrePost | 1 | 17 | 21.21 | **<.001** | 0.555 |
| Stimulus | 1 | 17 | 35.98 | **<.001** | 0.679 |
| Block | 1 | 17 | 92.02 | **<.001** | 0.844 |
| Intervention x PrePost | 1 | 17 | 0.75 | 0.399 | 0.042 |
| Intervention x Stimulus | 1 | 17 | 0.34 | 0.569 | 0.019 |
| PrePost x Stimulus | 1 | 17 | 11.84 | **0.003** | 0.41 |
| Intervention x Block | 1 | 17 | 0 | 0.983 | <.001 |
| PrePost x Block | 1 | 17 | 0.43 | 0.523 | 0.024 |
| Stimulus x Block | 1 | 17 | 64.89 | **<.001** | 0.792 |
| Intervention x PrePost x Stimulus | 1 | 17 | 0.45 | 0.513 | 0.026 |
| Intervention x PrePost x Block | 1 | 17 | 0.2 | 0.663 | 0.011 |
| Intervention x Stimulus x Block | 1 | 17 | 1.57 | 0.227 | 0.085 |
| PrePost x Stimulus x Block | 1 | 17 | 2.28 | 0.149 | 0.118 |
| Intervention x PrePost x Stimulus x Block | 1 | 17 | 0.03 | 0.854 | 0.002 |

#### Subjective ratings

*ANOVA of valence ratings to conditioned stimuli (Intervention: Sleep/Wake; PrePost: Before/After; Stimulus: CS+/CS-; Block: 0%, 50%, 100% reinforcement rate blocks)*

Table S11 – ANOVA of valence ratings to conditioned stimuli

| Predictor | $df_{Num}$ | $df_{Den}$ | $F$ | $p$ | $\eta^2$ |
|---|---|---|---|---|---|
| Intervention | 1 | 17 | 1.54 | 0.231 | 0.083 |
| PrePost | 1 | 17 | 0.67 | 0.426 | 0.038 |
| Stimulus | 1 | 17 | 14.87 | **0.001** | 0.467 |
| Block | 1.76 | 29.87 | 3.19 | 0.061 | 0.158 |
| Intervention x PrePost | 1 | 17 | 1.57 | 0.227 | 0.085 |
| Intervention x Stimulus | 1 | 17 | 0.05 | 0.821 | 0.003 |
| PrePost x Stimulus | 1 | 17 | 0.03 | 0.855 | 0.002 |
| Intervention x Block | 1.70 | 28.83 | 0.58 | 0.537 | 0.033 |
| PrePost x Block | 1.87 | 31.76 | 0.86 | 0.428 | 0.048 |
| Stimulus x Block | 1.23 | 20.98 | 4.61 | **0.037** | 0.213 |
| Intervention x PrePost x Stimulus | 1 | 17 | 0.01 | 0.938 | <.001 |
| Intervention x PrePost x Block | 1.56 | 26.45 | 0.31 | 0.680 | 0.018 |
| Intervention x Stimulus x Block | 1.98 | 33.69 | 0.48 | 0.622 | 0.027 |
| PrePost x Stimulus x Block | 1.99 | 33.88 | 0.72 | 0.492 | 0.041 |
| Intervention x PrePost x Stimulus x Block | 1.33 | 22.66 | 0.06 | 0.869 | 0.004 |

*ANOVA of valence ratings to conditioned stimuli (Intervention: Sleep/Wake; PrePost: Before/After; Stimulus: CS+/CS-; Block: 50%, 100% reinforcement rate blocks)*

Table S12 – ANOVA of valence ratings to conditioned stimuli

| Predictor | $df_{Num}$ | $df_{Den}$ | $F$ | $p$ | $\eta^2$ |
|---|---|---|---|---|---|
| Intervention | 1 | 17 | 1.82 | 0.195 | 0.097 |
| PrePost | 1 | 17 | 0.95 | 0.342 | 0.053 |
| Stimulus | 1 | 17 | 14.23 | **0.002** | 0.456 |
| Block | 1 | 17 | 0.35 | 0.559 | 0.020 |
| Intervention x PrePost | 1 | 17 | 1.34 | 0.262 | 0.073 |
| Intervention x Stimulus | 1 | 17 | 0 | >.999 | <.001 |
| PrePost x Stimulus | 1 | 17 | 0.5 | 0.489 | 0.029 |
| Intervention x Block | 1 | 17 | 0.57 | 0.461 | 0.032 |
| PrePost x Block | 1 | 17 | 0.94 | 0.346 | 0.052 |
| Stimulus x Block | 1 | 17 | 0.15 | 0.699 | 0.009 |
| Intervention x PrePost x Stimulus | 1 | 17 | 0.03 | 0.858 | 0.002 |
| Intervention x PrePost x Block | 1 | 17 | 0.02 | 0.893 | 0.001 |
| Intervention x Stimulus x Block | 1 | 17 | 0.03 | 0.872 | 0.002 |
| PrePost x Stimulus x Block | 1 | 17 | 0.13 | 0.719 | 0.008 |
| Intervention x PrePost x Stimulus x Block | 1 | 17 | 0.09 | 0.774 | 0.005 |

*ANOVA of arousal ratings to conditioned stimuli (Intervention: Sleep/Wake; PrePost: Before/After; Stimulus: CS+/CS-; Block: 0%, 50%, 100% reinforcement rate blocks)*

Table S13 – ANOVA of arousal ratings to conditioned stimuli

| Predictor | $df_{Num}$ | $df_{Den}$ | $F$ | $p$ | $\eta^2$ |
|---|---|---|---|---|---|
| Intervention | 1 | 17 | 0.72 | 0.408 | 0.041 |
| PrePost | 1 | 17 | 0.41 | 0.532 | 0.023 |
| Stimulus | 1 | 17 | 1.38 | 0.255 | 0.075 |
| Block | 1.33 | 22.68 | 3.50 | 0.064 | 0.171 |
| Intervention x PrePost | 1 | 17 | 0.02 | 0.900 | <.001 |
| Intervention x Stimulus | 1 | 17 | 0.01 | 0.924 | <.001 |
| PrePost x Stimulus | 1 | 17 | 1.31 | 0.269 | 0.071 |
| Intervention x Block | 1.54 | 26.25 | 0.67 | 0.484 | 0.038 |
| PrePost x Block | 1.83 | 31.17 | 1.15 | 0.327 | 0.063 |
| Stimulus x Block | 1.64 | 27.8 | 2.35 | 0.122 | 0.122 |
| Intervention x PrePost x Stimulus | 1 | 17 | 0.48 | 0.499 | 0.027 |
| Intervention x PrePost x Block | 1.70 | 28.82 | 0.93 | 0.391 | 0.052 |
| Intervention x Stimulus x Block | 1.97 | 33.54 | 1.6 | 0.218 | 0.086 |
| PrePost x Stimulus x Block | 1.33 | 22.69 | 0.43 | 0.574 | 0.025 |
| Intervention x PrePost x Stimulus x Block | 1.90 | 32.24 | 0.66 | 0.517 | 0.037 |

*ANOVA of arousal ratings to conditioned stimuli (Intervention: Sleep/Wake; PrePost: Before/After; Stimulus: CS+/CS-; Block: 50%, 100% reinforcement rate blocks)*

Table S14 – ANOVA of arousal ratings to conditioned stimuli

| Predictor | $df_{Num}$ | $df_{Den}$ | $F$ | $p$ | $\eta^2$ |
|---|---|---|---|---|---|
| Intervention | 1 | 17 | 0.78 | 0.388 | 0.044 |
| PrePost | 1 | 17 | 0 | 0.963 | <.001 |
| Stimulus | 1 | 17 | 2.19 | 0.158 | 0.114 |
| Block | 1 | 17 | 2.05 | 0.171 | 0.107 |
| Intervention x PrePost | 1 | 17 | 0.2 | 0.658 | 0.012 |
| Intervention x Stimulus | 1 | 17 | 0 | 0.969 | <.001 |
| PrePost x Stimulus | 1 | 17 | 1.19 | 0.290 | 0.066 |
| Intervention x Block | 1 | 17 | 1 | 0.331 | 0.056 |
| PrePost x Block | 1 | 17 | 0.74 | 0.401 | 0.042 |
| Stimulus x Block | 1 | 17 | 0.11 | 0.745 | 0.006 |
| Intervention x PrePost x Stimulus | 1 | 17 | 0.68 | 0.420 | 0.039 |
| Intervention x PrePost x Block | 1 | 17 | 0.06 | 0.816 | 0.003 |
| Intervention x Stimulus x Block | 1 | 17 | 3.58 | 0.076 | 0.174 |
| PrePost x Stimulus x Block | 1 | 17 | 0.01 | 0.919 | <.001 |
| Intervention x PrePost x Stimulus x Block | 1 | 17 | 0.47 | 0.501 | 0.027 |

*ANOVA of valence and arousal ratings to unconditioned stimuli (Intervention: Sleep/Wake; PrePost: Before/After; Stimulus: CS+/CS-; Block: 50%, 100% reinforcement rate blocks)*

Table S15 – ANOVA of valence ratings to unconditioned stimuli

| Predictor | $df_{Num}$ | $df_{Den}$ | *F* | *p* | $\eta^2$ |
|---|---|---|---|---|---|
| Intervention | 1 | 17 | 0.37 | 0.551 | 0.021 |
| PrePost | 1 | 17 | 0.02 | 0.899 | <.001 |
| Stimulus | 1 | 17 | 122.92 | **<.001** | 0.878 |
| Block | 1 | 17 | 3.21 | 0.091 | 0.159 |
| Intervention x PrePost | 1 | 17 | 0.23 | 0.639 | 0.013 |
| Intervention x Stimulus | 1 | 17 | 4.05 | 0.060 | 0.192 |
| PrePost x Stimulus | 1 | 17 | 2.29 | 0.149 | 0.119 |
| Intervention x Block | 1 | 17 | 1.19 | 0.291 | 0.065 |
| PrePost x Block | 1 | 17 | 1.48 | 0.240 | 0.080 |
| Stimulus x Block | 1 | 17 | 0.61 | 0.444 | 0.035 |
| Intervention x PrePost x Stimulus | 1 | 17 | 1.21 | 0.286 | 0.067 |
| Intervention x PrePost x Block | 1 | 17 | 0.01 | 0.908 | <.001 |
| Intervention x Stimulus x Block | 1 | 17 | 0.18 | 0.681 | 0.010 |
| PrePost x Stimulus x Block | 1 | 17 | 0.04 | 0.842 | 0.002 |
| Intervention x PrePost x Stimulus x Block | 1 | 17 | 0.01 | 0.940 | <.001 |

Table S16 – ANOVA of arousal ratings to unconditioned stimuli

| Predictor | $df_{Num}$ | $df_{Den}$ | *F* | *p* | $\eta^2$ |
|---|---|---|---|---|---|
| Intervention | 1 | 17 | 0.49 | 0.493 | 0.028 |
| PrePost | 1 | 17 | 6.94 | **0.017** | 0.29 |
| Stimulus | 1 | 17 | 28.87 | **<.001** | 0.629 |
| Block | 1 | 17 | 2.97 | 0.103 | 0.149 |
| Intervention x PrePost | 1 | 17 | 0.91 | 0.353 | 0.051 |
| Intervention x Stimulus | 1 | 17 | 1.26 | 0.278 | 0.069 |
| PrePost x Stimulus | 1 | 17 | 0.18 | 0.678 | 0.01 |
| Intervention x Block | 1 | 17 | 1.67 | 0.214 | 0.089 |
| PrePost x Block | 1 | 17 | 0.04 | 0.848 | 0.002 |
| Stimulus x Block | 1 | 17 | 0.63 | 0.437 | 0.036 |
| Intervention x PrePost x Stimulus | 1 | 17 | 1.16 | 0.297 | 0.064 |
| Intervention x PrePost x Block | 1 | 17 | 0.57 | 0.46 | 0.032 |
| Intervention x Stimulus x Block | 1 | 17 | 0.18 | 0.677 | 0.01 |
| PrePost x Stimulus x Block | 1 | 17 | 0.15 | 0.705 | 0.009 |
| Intervention x PrePost x Stimulus x Block | 1 | 17 | 0.06 | 0.811 | 0.003 |
